# Repurposing the CRISPR-Cas9 System for Targeted Chromatin O-linked β-N-acetylglucosamine Editing

**DOI:** 10.1101/2022.10.27.514044

**Authors:** Matthew P. Parker, Wagner B. Dias, Will Brautman, Nick Lowe, Halyna Fedosyuk, Kenneth R. Peterson, Chad Slawson

## Abstract

Eukaryotic gene transcription is controlled by many proteins, including the basal transcription machinery, epigenetic chromatin remodeling complexes, and transcription cofactors. Chromatin and genome-mapping consortia identified *O*-linked β-*N*-acetylglucosamine (O-GlcNAc) as an abundant chromatin post-translational modification involved in numerous transcriptional processes, including RNA polymerase function, epigenetic dynamics, and transcription factor activity. Thus, O-GlcNAc regulation of *cis*-regulatory elements is essential for proper gene expression. O-GlcNAc is a single N-acetylglucosamine sugar attached to serine or threonine residues in nuclear, cytoplasmic, or mitochondrial proteins. Two enzymes cycle O-GlcNAc on or off protein; O-GlcNAc transferase (OGT) adds the modification, and O-GlcNAcase (OGA) removes it. O-GlcNAcylation responds to inputs from multiple metabolic and stress pathways including glucose, amino acid, fatty acid, and nucleotide metabolism. Therefore, O-GlcNAc acts as a sensor of cellular homeostasis able to link environmental conditions with gene transcription; however, decoding the precise function of millions of O-GlcNAc regulated elements remains challenging. Technologies to readily manipulate O-GlcNAcylation at specific *cis*-regulatory elements for functional analysis without pleiotropic consequences are lacking. We have employed novel CRISPR-based gene targeting tools to probe the function of O-GlcNAc regulated *cis*-elements. First, we developed a programmable CRISPR-Cas9-based targeting system. This was accomplished by fusing a catalytically dead Cas9 (dCas9) to O-GlcNAc transferase (OGT) or O-GlcNAcase (OGA), which allows for highly specific O-GlcNAc manipulation at chromatin *cis*-regulatory elements. Previously, we demonstrated that O-GlcNAc plays a role in regulating human ^A^***γ***-globin gene expression by regulating CHD4 function and the formation of the NuRD (Nucleosome Remodeling and Deacetylase) complex at the -566 GATA repressor-binding site. Thus, as a proof of principle and to further explore the function of O-GlcNAc in ***γ***-globin gene transcription, we targeted both dCas9-OGT and -OGA fusion proteins to the ^A^***γ***-globin gene promoter. When dCas9-OGT or dCas9-OGA was targeted to the -566 GATA silencer site of the ^A^***γ***-globin promoter, gene expression decreased or increased, respectively. This data strongly correlates with our previous findings and implicates O-GlcNAc cycling in ***γ***-globin gene regulation. Importantly, this method can be employed to investigate O-GlcNAc events known to exist within the eukaryotic genome in a highly specific manner. Together, this tool will be fundamental in elucidating the function of O-GlcNAc in gene transcription.

## Introduction

O-GlcNAcylation (O-GlcNAc) is a ubiquitous post-translational modification involving the attachment of a single N-acetylglucosamine sugar to serine or threonine residues of nuclear, cytoplasmic, and mitochondrial proteins (1). O-GlcNAc modulates protein activity, localization, and stability through the coordinated action of O-GlcNAc transferase (OGT) and O-GlcNAcase (OGA), which dynamically cycle the modification on and off proteins, respectively (2). All aspects of metazoan transcription are sensitive to O-GlcNAc, including RNA polymerase function, epigenetic dynamics, and transcription factor activity (3). UDP-GlcNAc, the substrate for OGT and the end product of the <u>H</u>exosamine <u>B</u>iosynthetic <u>P</u>athway (HBP), is sensitive to fluctuations in amino acid, fatty acid, nucleotide, and glucose metabolism (4). Thus, O-GlcNAcylation of the transcriptional machinery changes gene expression profiles in response to nutrient flux and stress (5). Furthermore, O-GlcNAc is key to organizing large transcriptional complexes at *cis*-regulatory elements. OGT and OGA interact with many epigenetic “writers” and “erasers” of the histone code (6). For example, OGT interacts with chromatin remodeling complexes such as the SET1/COMPASS complex, polycomb complex (PRC2), mSin3a complex, and nucleosome remodeling deacetylase (NuRD) complex, all of which are essential for proper gene regulation (3,7,8). O-GlcNAc is also part of the histone code. Numerous O-GlcNAcylation sites have been identified on histones, including several at the histone-DNA interface. However, the function of the modification at this interface is unclear (1). Additionally, O-GlcNAcylation modulates the ubiquitination and methylation of histones (9,10). O-GlcNAcylation is involved in the function of the basal transcription machinery (11). For example, assembly of the pre-initiation complex (PIC) requires O-GlcNAcylation of the C-terminal domain (CTD) of RNA polymerase II (RNAP II). As transcription proceeds, OGA removes O-GlcNAc from the CTD of RNAP II prior to CTD phosphorylation and mRNA elongation (6,12–14). In addition to RNAP II, mass spectroscopy data has identified 32 additional RNAP II transcription cycling factors that are O-GlcNAcylated (15). Recently, mutations of the O-GlcNAc site on the TATA-box protein led to large-scale rearrangement of gene expression patterns at predominantly metabolic genes suggesting that O-GlcNAc controls the PIC (11). However, the number of promoters that use this form of gene regulation is unclear, as only a small number of promoters have been extensive investigated thus far.

Understanding the roles OGT, OGA, and O-GlcNAcylation play in gene transcription is difficult for many reasons. Both enzymes are essential for cellular growth, development, and in most cases, survival (16). In rapidly dividing tissue, the loss of OGT or OGA is lethal, making knockout studies problematic (17). Further complicating O-GlcNAc experimental studies is the fact that the expression of OGT and OGA are transcriptionally linked (18,19); that is, cells maintain a set level of O-GlcNAcylation. Changes in cellular levels of O-GlcNAcylation lead to rapid changes in both OGT and OGA protein levels as a mechanism for maintaining O-GlcNAc homeostasis. Experimental manipulations that alter the protein level of one enzyme lead to compensatory changes in protein levels of the other enzyme, thereby limiting the amount of knowledge that can be gained using these approaches (19–21). Moreover, one of the most daunting issues when trying to determine the role of O-GlcNAc in transcription is the overwhelming number of cellular processes sensitive to O-GlcNAc manipulation. To date, more than 5,000 O-GlcNAcylated proteins have been identified, complicating the ability to study specific O-GlcNAc events in the cell. Genetic or pharmacological manipulation of OGT and OGA affects all these proteins. Hence, pleiotropic effects make the results of these studies difficult to interpret. Additionally, because the modification is found on serine and threonine residues, it has extensive crosstalk with phosphorylation. This is problematic, as serine or threonine to alanine substitutions intended to block O-GlcNAcylation also block phosphorylation at these residues, confounding data interpretation. Understanding the function of O-GlcNAc in transcription required the development of new technologies. Our approach exploits the genome targeting capabilities of the CRISPR/Cas9 system to address questions surrounding the role of O-GlcNAc in gene transcription.

Advances in CRISPR/Cas9 technology allow researchers to target proteins of interest to specific *cis*-regulatory regions such as promoters, enhancers, and silencers. The Streptococcus pyogenes type II CRISPR/Cas9 system is a versatile tool for genome engineering and functional genomic studies (22–25). The RNA-guided Cas9 endonuclease protein can be concentrated at specific genomic loci via complementarity between an engineered single-guide RNA (sgRNA) and the target DNA sequence (26–28). Abolishing the enzymatic activity of Cas9 endonuclease is accomplished by mutations to the RuvC catalytic domain (D10A) and HNH catalytic domain (H840A), which generates a catalytically dead Cas9 (dCas9) protein (27). Numerous synthetic transcriptional regulators fused dCas9 such as the catalytic domain of p300 acetyltransferase (dCas9-p300) or the Krüppel-associated box (KRAB) repressor domain have been used to modulate gene expression (24,29). Thus, we hypothesize that similar systems can be developed to study the function of OGT and OGA at distinct *cis*-regulatory elements (**Figure 1**).

**Figure 1:**
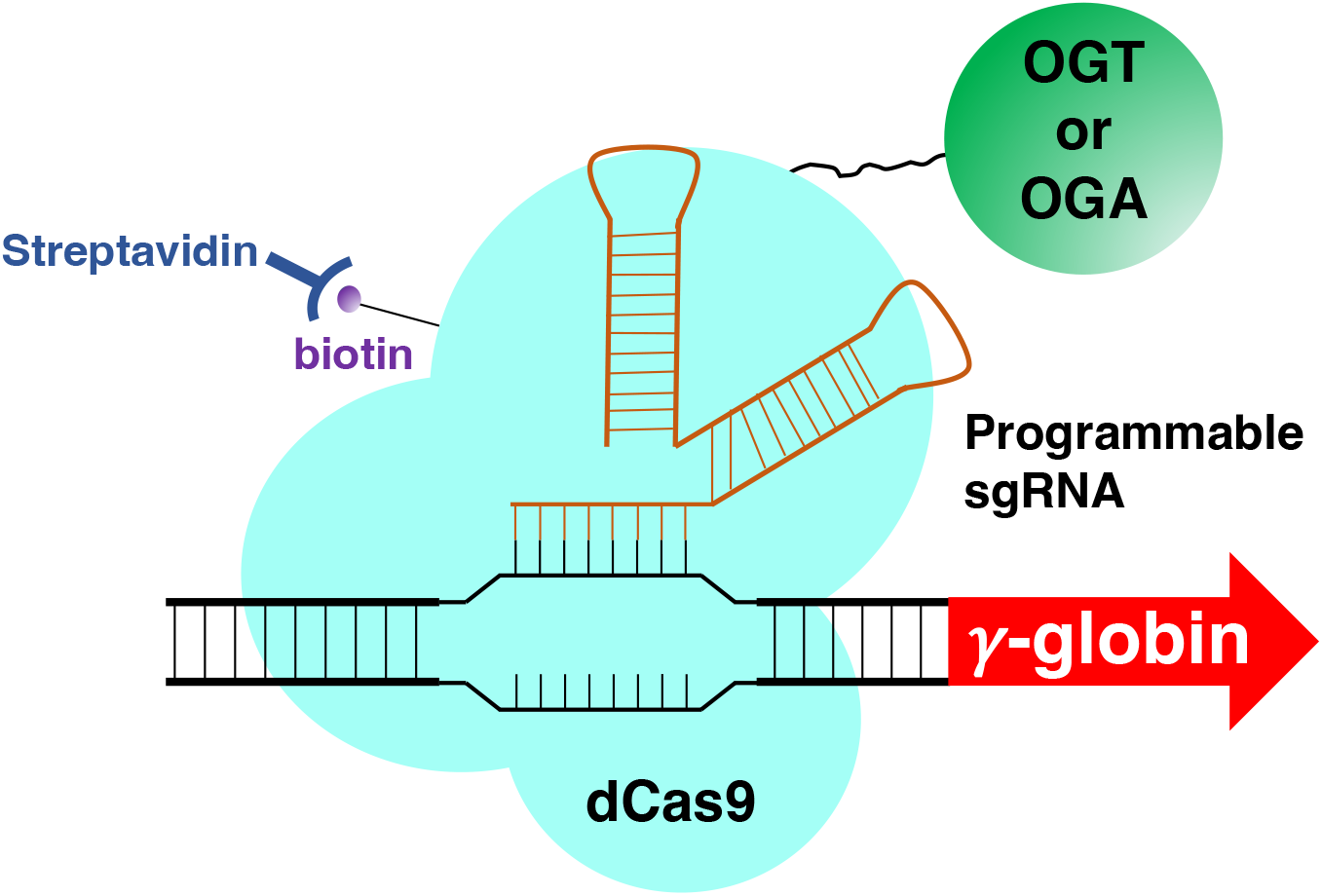
Schematic diagram of dCas9 OGT or OGA prototype construct. OGT, OGA, or catalytically dead versions of OGT and OGA were fused to a catalytically dead Cas9 (dCas9). The dCas9 contains a FLAG tag and an AviTag. Specific guide (sg) RNAs can deliver the fusion proteins to specific *cis*-regulatory elements in the ^A^γ-globin promoter.

Previously, we demonstrated that O-GlcNAc plays a role in regulating human γ-globin gene transcription (30,31). Increasing fetal hemoglobin (HbF) via activation of γ-globin chain synthesis is widely accepted as the most effective treatment for Sickle Cell Disease (SCD) and certain types of β-thalassemias. O-GlcNAcylation modulates the formation of a GATA-1-FOG-1-NuRD repressor complex that binds the −566/567 GATA-1 site of the ^A^γ- and ^G^γ-globin promoters, respectively, when γ-globin genes are silent. O-GlcNAcylation of Chromodomain Helicase DNA Binding Protein 4 (CHD4; previously known as Mi2β), a critical component of the NuRD complex, is necessary to form this repressor complex. OGA inhibition using Thiamet-G (TMG) prevents the removal of O-GlcNAc from CHD4 and maintains repression of the γ-globin genes (30,31). At the γ-globin genes, the NuRD complex is composed of GATA Zinc Finger Domain Containing 2A (GATAD2A), Histone Deacetylase 2 (HDAC2), Retinoblastoma-binding protein 4 (RBBP4), and methyl cytosine-guanosine (cpG)-binding domain protein 2 (MBD2), and metastasis-associated factor (MTA) family member 2 (MTA2) (32). Recent studies have identified O-GlcNAc on every protein of the NuRD complex; however, the function of these modifications remains unknown (33,34). As a proof of principle, and to explore the function of O-GlcNAc in ^A^γ-globin gene transcription, we targeted both dCas9-OGT and OGA fusion proteins to DNA sequences flanking the −566 GATA site of the ^A^γ-globin gene promoter (**Figure 1**). Gene expression analysis by RT-qPCR revealed increased γ-globin gene expression when dCas9-OGA was targeted to this area and decreased γ-globin gene expression when dCas9-OGT was targeted to identical sequences, further supporting our previous data implicating O-GlcNAcylation in ^A^γ-globin gene regulation. These data demonstrate the utility of targeting OGT and OGA to promoters and shows the potential for the dCAS9-OGT and OGA constructs as tools to probe the role of these proteins in transcriptional complexes at *cis*-regulatory elements within the genome.

## Results

### Creation of plasmids for expression of OGA, OGT, dOGA, and dOGT fusion proteins and generation of stable monoclonal cell lines

To achieve targeted editing of chromatin via O-GlcNAcylation, two expression cassettes were engineered. Each plasmid construct was assembled in the same linear arrangement (5’ to 3’): 1) pUC19 backbone with an elongation factor 1-alpha (EF1α) promoter; 2) OGA (**Figure 2a**) or OGT (**Figure 2c**); 3) a *Not*I restriction site and a flexible alanine linker; 4) dCas9; and 5) a FLAG-tag and biotin-acceptor-site fused back to the pUC19 plasmid backbone. We confirmed that the products were correctly assembled with restriction enzyme-digests and Sanger di-deoxy sequencing.

**Figure 2:**
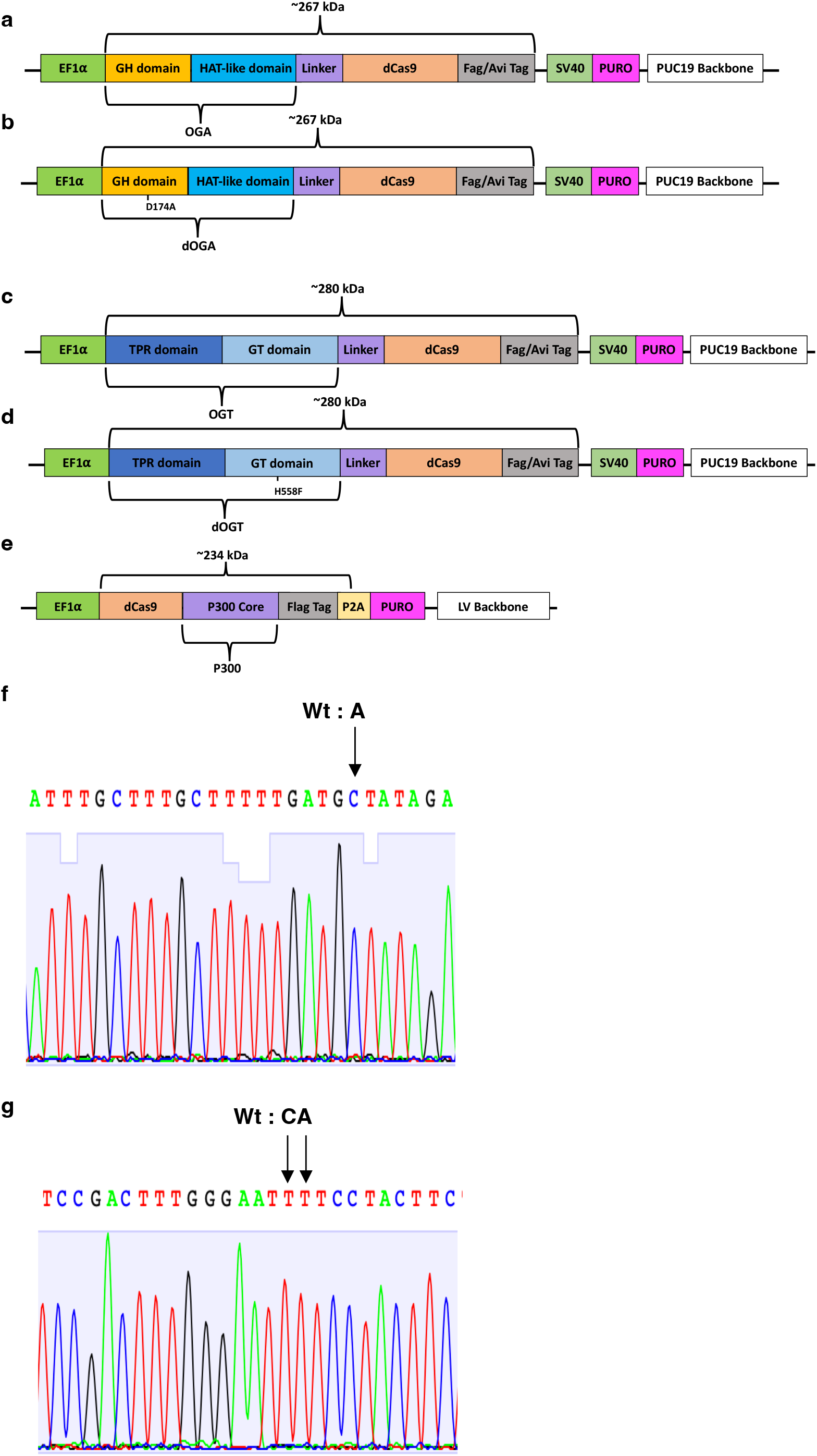
Schematic of dCAS9 fusion plasmids. (**a**) OGA-dCas9 plasmid. (**b**) dOGA-dCas9 plasmid. (**c**) OGT-dCas9 plasmid. (**d**) dOGT-dCas9 plasmid. (**e**) dCas9-P300 plasmid. (**f**) Sanger dideoxy sequencing data confirming site-directed mutagenesis of OGA-dCas9 plasmid to produce dOGA-dCas9. (**g**) Sanger dideoxy sequencing data confirming site-directed mutagenesis of OGT-dCas9 plasmid to produce dOGT-dCas9.

Independent of catalytic function, OGA and OGT can also function as bridge proteins or act as scaffolds for binding to or recruiting other transcriptional proteins that regulate gene expression (35–37). To interrogate how the enzymatic activity of OGA or OGT influences proteins associated with *cis*-regulatory elements and gene expression, we used our OGA-dCas9 and OGT-dCas9 plasmid DNAs to generate several experimental controls. These controls consisted of catalytically dead OGA (dOGA)- or OGT (dOGT)-dCas9 expression cassettes. To generate them, we perform site-directed mutagenesis on OGA or OGT. For OGA, we introduced a D174>A mutation (**Figure 2b and Figure 2f**), and for OGT, we introduced a H558>F mutation (**Figure 2d and Figure 2g)**. These mutations produce stable mutant proteins that abolish the catalytic activity of the enzymes (38,39). The final products were confirmed with restriction enzyme-digests and Sanger di-deoxy sequencing (**Figure 2a-d**). Additionally, a dCas9-P300 expression construct was used as a positive control for gene activation in our experiments (**Figure 2e**); this construct has been described elsewhere (40).

All constructs were linearized and introduced into K562 cells, a well-characterized human erythro-leukemia cell line, via electroporation. After puromycin selection, limiting dilution was employed to generate monoclonal cell lines stably expressing the fusion protein transgene. DNA PCR genotyping and Western blots confirmed the expression of all fusion protein constructs (**Figure 3a-c**). Monoclonal cell lines that contained similar gene cassette copy numbers and fusion protein expression levels were selected for subsequent experiments.

**Figure 3:**
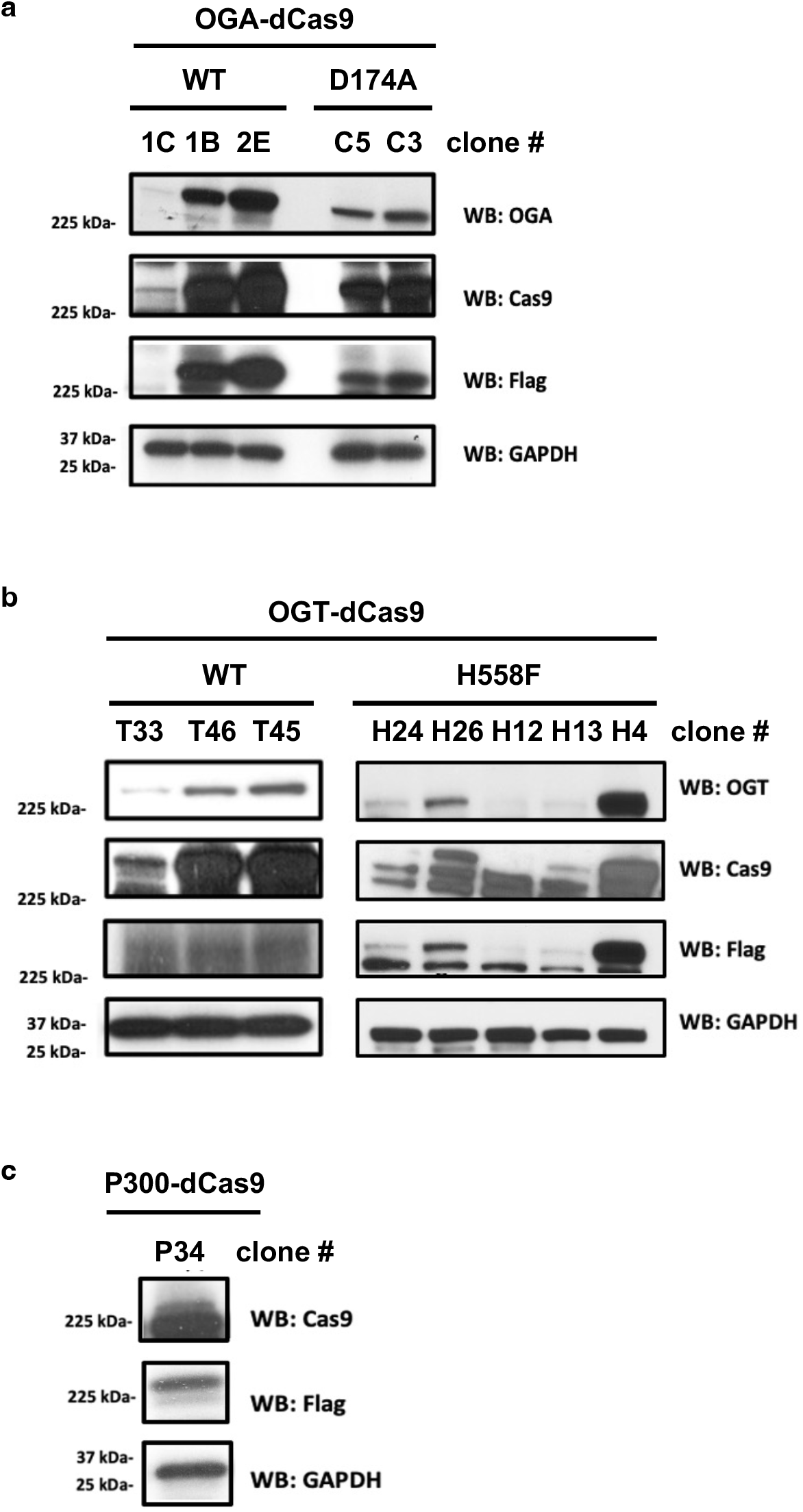
Fusion expression in monoclonal K562 cell lines. 50μg of whole cell lysates was used for electrophoresis and subjected to Western blot analysis. Western blots were queried with antibodies specific to Cas9, OGT, OGA, and FLAG. GAPDH was utilized as a loading control. (**a**) Stable monoclonal cell lines expressing different levels of OGA-dCas9 and dOGA-dCas9 fusion proteins (**b**) Stable monoclonal cell lines expressing different levels of OGT-dCas9 fusion proteins. (**c**) Stable monoclonal cell line expressing dCas9-P300 fusion protein.

### Synthesis of sgRNA plasmids and generation of sgRNA stable cell lines

There is a wealth of genomic information in the ENCODE database for K562 cells that helped us identify DNA *cis*-regulatory sequences to validate our new CRISPR/dCas9 technology. We elected to utilize *cis*-regulatory elements that bind OGT within the well-characterized human β-globin locus. The human β-globin locus consists of five functional β-like globin genes, each of which serves as the β-like chain in the hemoglobin molecule during different stages of development. The β-globin locus control region (LCR) is a super-enhancer composed of five DNAse I hypersensitive sites (5’HS1-5) and is positioned upstream of the five β-like globin genes (HBE1, HBG1, HBG2, HBD, HBB) (41). The LCR orchestrates transcription of the downstream β-like globin genes at different times throughout development in erythroid cells (40). K562 cells have an intact LCR and express basal levels of HBE1 (ε-globin), HBG1(^G^γ-globin), and HBG2 (^A^γ-globin) (42). We, and others, have identified OGT and OGA interacting sites in several of the β-like globin gene promoters and in the LCR (31,34,43). Many sgRNA sequences have already been generated and validated for this locus due to its pathophysiological relevance in human diseases, such as SCD and certain types of β-thalassemia. From our previously published data, we know the transcription factor GATA-1 forms a repressor complex containing FOG-1, OGT, OGA, and CHD4 at the −566 or −567 GATA-1 binding sites of the γ-globin promoters (30,31). To validate our O-GlcNAc targeting systems, we chose to interrogate the ^A^γ-globin gene promoter, with emphasis on the −566 GATA-1 binding site.

Utilizing the ENCODE database, we generated a small library of nine sgRNAs that did not overlap with any known transcription factor binding sites (**Figure 4a**) with two guides that flank the −566 GATA-1 binding site. Three non-targeting sgRNAs would serve as controls (**Figure 4a and b**). The sgRNA sequences were ligated into the LentiCRISPRv2 hygro plasmid, which drives expression of the sgRNA from an elongation factor 1-alpha (EF1α) promoter. The plasmids were linearized and introduced into our stable dCAS9-OGT/OGA monoclonal K562 cell lines. After hygromycin selection, the cells were analyzed for expression of ^A^γ-globin mRNA using RT-qPCR.

**Figure 4:**
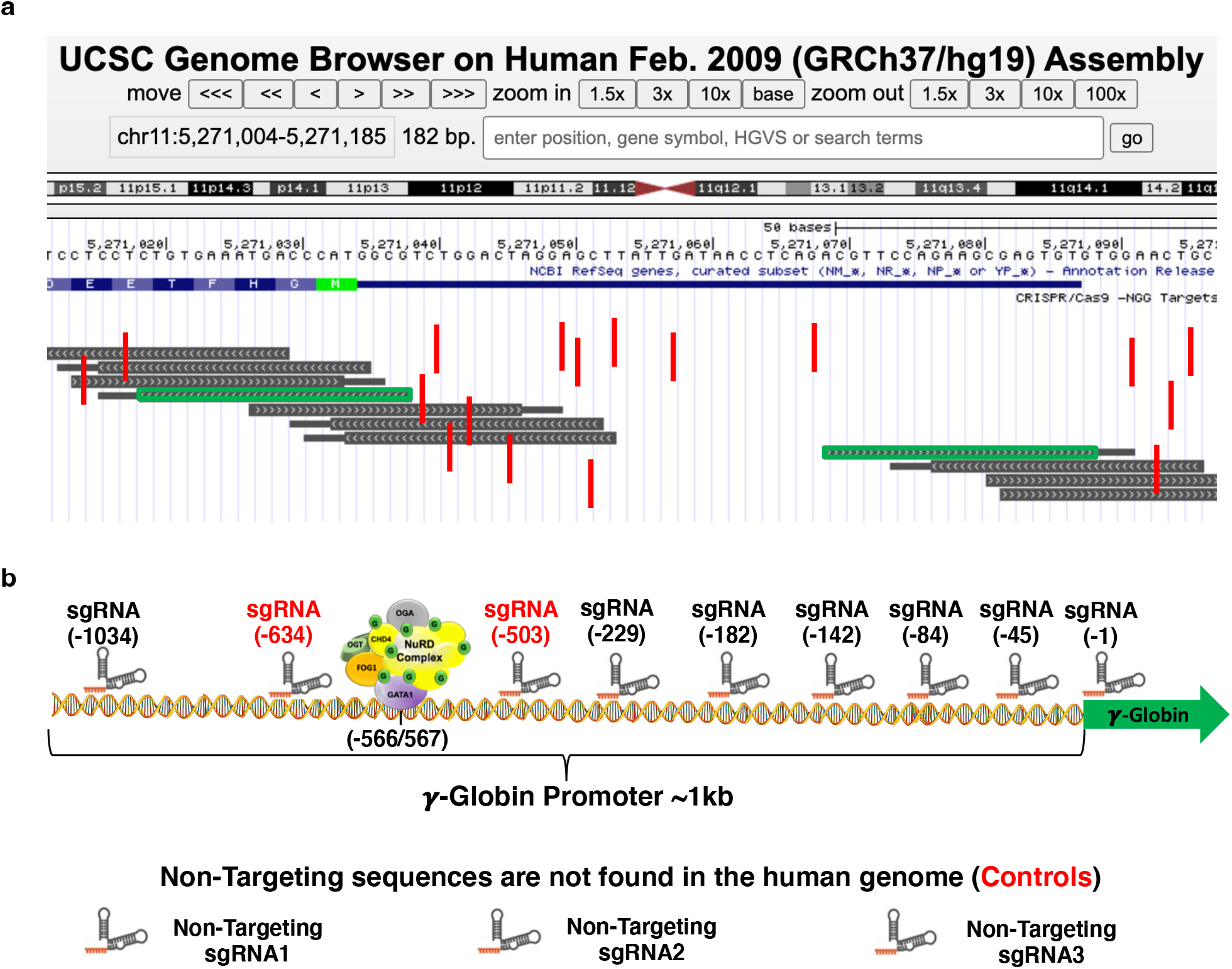
sgRNA targets in the ^A^γ-globin promoter of the human β-globin locus. (**a**) ENCODE data set (Feb. 2009 (GRCh37/hg19) Assembly) was used to generate a sgRNA library for the ^A^γ-globin promoter. sgRNAs that did not disrupt known transcription factor binding site (vertical red bars) were selected (green rectangles) for use in targeting dCAS9 fusion proteins. (**b**) sgRNAs selected for incorporation into plasmid constructs for use in ^A^γ-globin *cis*-regulatory element targeting experiments are shown. Nine sgRNA sequences were selected. Locations of the sgRNAs are indicated. The numbers represent position relative to the ^A^γ-globin gene transcriptional start site. Three non-targeting sgRNAs were selected and used as controls in subsequent experiments. Two sgRNAs (red) that flank the ^A^γ-globin -566GATA-1-FOG-1-OGT-OGA-NuRD repressor complex binding site are shown.

### dCas9-P300 targeting increases ^A^γ-Globin mRNA expression

As more information has become available for CRISPR/Cas9 methodologies, it has become clear that not all sgRNAs are equal functionally. To achieve successful targeting of Cas9 protein, sgRNAs must form the correct 3-dimensional structure (44,45). It is difficult to predict how the 20-nucleotide targeting sequence introduced into the sgRNA molecule will affect its overall structure. To ensure that our sgRNA cassettes produced functional sgRNAs, we introduced them into our dCas9-P300 stable cell line. The P300 histone acetyltransferase (HAT) core domain produced by this construct acetylates histones when targeted to genomic sequences. This leads to gene activation (24,46,47). RT-qPCR data demonstrated that the dCas9-P300 fusion protein activated transcription of the ^A^γ-globin gene when delivered to the proximal promoter (**Figure 5**). The results of these experiments confirmed that all sgRNA cassettes produced functional sgRNAs and validated previously published data (24,46,47).

**Figure 5:**
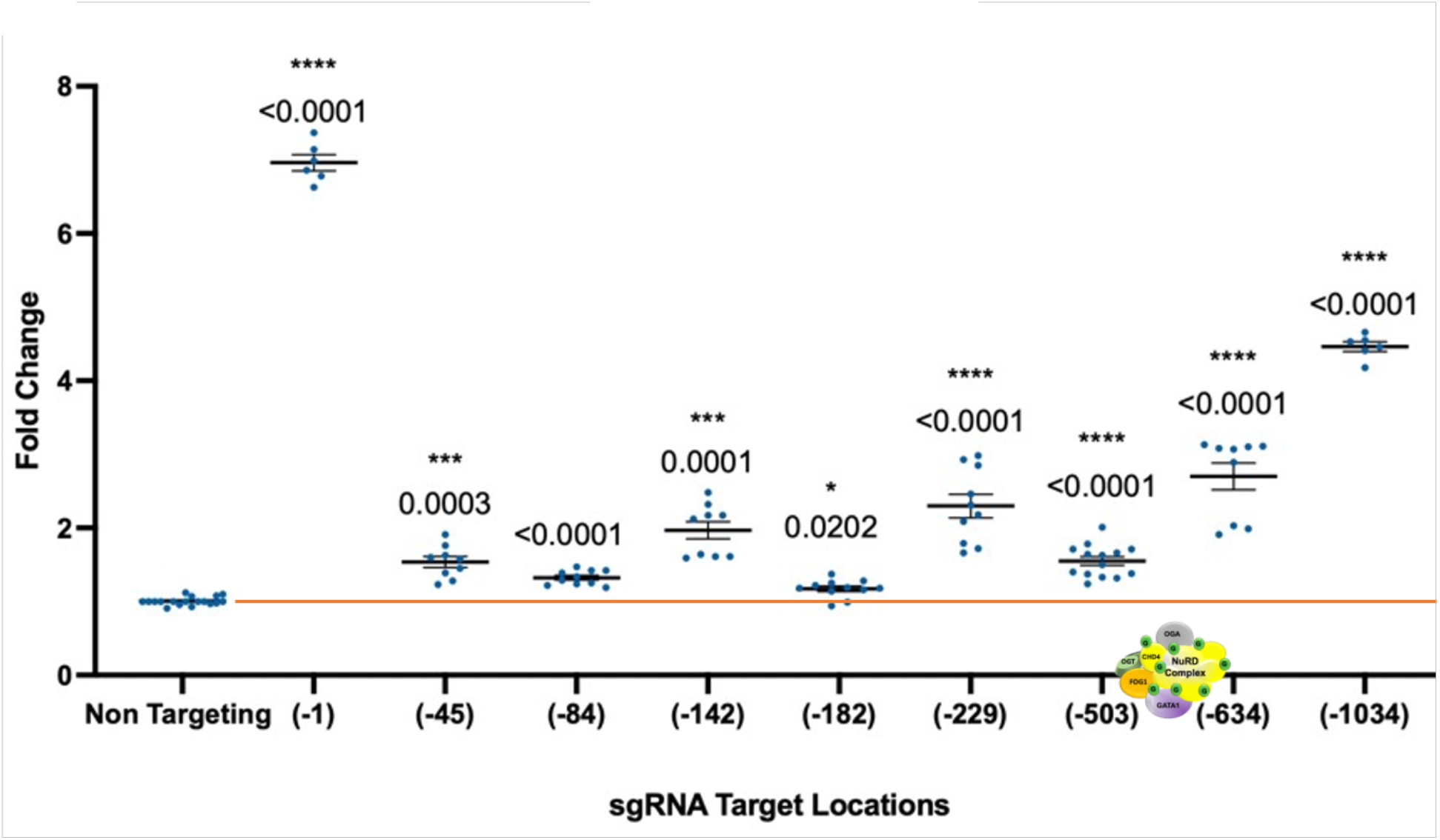
Confirmation of sgRNA function in a dCas9-P300 K562 monoclonal cell line. Relative mRNA expression of the ^A^γ-globin gene was determined by RT-qPCR. The dCas9-P300 fusion protein activates transcription (dots above the red line) of the ^A^γ-globin gene from the proximal promoter. Fold change was calculated using five housekeeping genes (ACTB, RNA18sN5, PKG1, GAPDH, and RPL13). All samples were normalized to non-targeting controls. Data generated in at least five independent experiments are presented as mean + or − standard error; the two-tailed Student t-test statistic was applied with p < 0.05 considered to be a significant difference. The location of the −566 GATA-1-FOG-1-NuRD repressor complex binding site is shown on the x-axis.

### OGA-dCas9 targeting increases γ-Globin mRNA expression

After confirming that all our sgRNA constructs produced functional sgRNAs, we used RT-qPCR to analyze ^A^γ-globin mRNA expression after targeting OGA-dCas9 and dOGA-dCas9 to the ^A^γ-globin promoter. The OGA-dCas9 fusion protein activated transcription of the ^A^γ-globin gene at all sgRNA target sequences compared to non-targeting controls (**Figure 6**). The mean fold increase in ^A^γ-globin expression ranged from ~1.36 to ~2.55. We observed the highest mean fold increases when OGA-dCas9 was targeted to sites flanking the −566 GATA-1-FOG-1-NuRD repressor binding site. The dOGA-dCas9 fusion protein also activated transcription of the ^A^γ-globin gene when targeted to multiple sites along the ^A^γ-globin promoter The mean fold increase ranged from ~1.52 to ~2.17, with the highest increase detected when dOGA-dCas9 was targeted to the −229 sgRNA target site.

**Figure 6:**
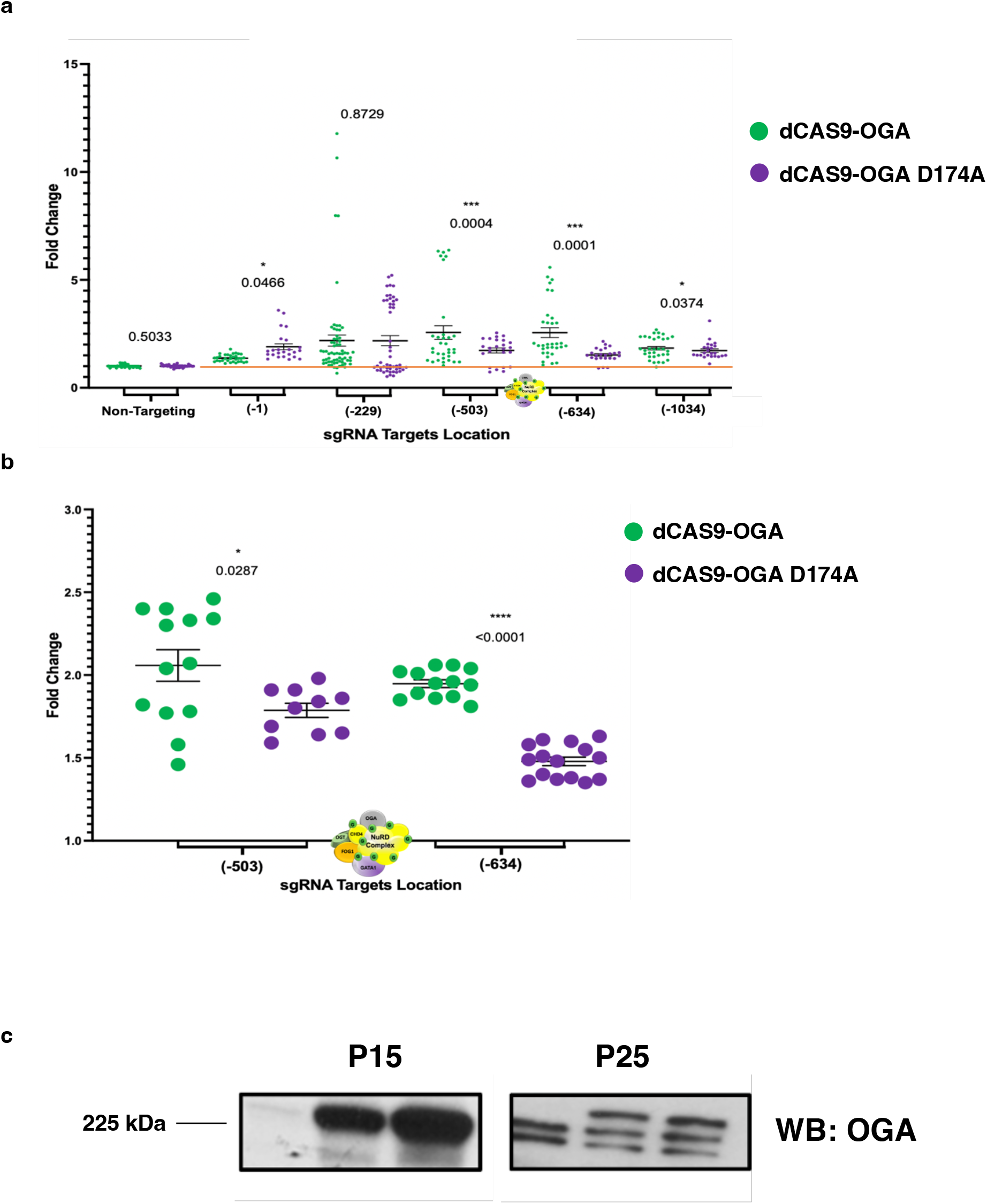
^A^γ-globin gene expression following targeting of OGA-dCas9 or dOGA-dCas9 fusion proteins to the gene promoter region. (**a**) Relative mRNA expression of the ^A^γ-globin gene was determined by RT-qPCR. The OGA-dCas9 fusion protein activates transcription (green dots above the red line) of the gene at all sgRNA target sequences compared to non-targeting controls. A similar outcome was noted with the dOGA-dCas9 fusion (**b**) OGA-dCas9 versus dOGA-dCas9 ^A^γ-globin mRNA expression at sgRNA target sites flanking the −566GATA-1-FOG-1-NuRD repressor binding site. OGA-dCas9 showed a statistically significant increases in γ-globin mRNA expression at both flanking sites. Fold change was calculated using five housekeeping genes (PPIA, RNA18sN5, PKG1, GAPDH, and RPLPO). All sample were normalized to non-targeting controls. Data generated in at least five independent experiments are presented as mean + or − standard error; the two-tailed Student t-test statistic was applied with p < 0.05 considered to be a significant difference. The location of the −566GATA-1-FOG-1-NuRD repressor binding site is shown on the x-axis. (**c**) The same OGA-dCas9 monoclonal cell line at different passages. Western blot analysis shows that cells from earlier passages express higher amounts of fusion proteins (top band).

When OGA-dCas9 was compared to dOGA-dCas9, OGA-dCas9 had statistically significant increases in γ-globin mRNA expression at three sgRNA target sites (−503, −634, and −1034). dOGA-dCas9 showed higher ^A^γ-globin mRNA expression at the −1 sgRNA target site when compared to OGA-dCas9. No statistically significant changes were observed between OGA-dCas9 and dOGA-dCas9 at the −229 sgRNA target site; however, both constructs had a mean fold increase of ~2.18 at that site (**Figure 6a**). We next focused on the two sgRNA sites flanking the −566 GATA-1-FOG-1-NuRD repressor binding site. OGA-dCas9 showed a statistically significant increase in ^A^γ-globin mRNA expression at both sites flanking the −566 GATA-1-FOG-1-NuRD repressor binding site (**Figure 6b)** compared to dOGA-dCas9.

Large fold changes were observed in some our experiments. Further investigation revealed that cells from earlier passages expressed higher amounts of our fusion protein constructs (**Figure 6c**). The formation of a functional ribonuclear Cas9 holoenzyme complex is stochastic; thus, increasing the cellular concentrations of both dCas9 fusion protein and sgRNA increases collisional frequencies and the likelihood of forming a functional complex. The ability of Cas9 proteins to find their target sequence is also stochastic in nature; thus, higher concentrations of Cas9 fusion protein also increases the collisional frequency with target sequences. This explains why we observed greater activation and repression with higher transgene expressing cells at low passages; thus, we only compared cell lines of approximately the same passage. Additionally, we used the GraphPad Prism 9 ROUT method to exclude statistical outliers.

### OGT-dCas9 targeting decreases γ-Globin mRNA expression

RT-qPCR was again used to analyze γ-globin mRNA expression after targeting OGT-dCas9 and dOGT-dCas9 to the ^A^γ-globin promoter. The OGT-dCas9 fusion protein had a significant repressive effect on ^A^γ-globin expression at all sgRNA target sequences compared to non-targeting controls. The dOGT-dCas9 fusion protein had both repressive (−1 and −45) and activating (−503 and −634) effects on ^A^γ-globin mRNA expression compared to non-targeting controls. Again, the highest repressive effects were observed when dOGT-dCas9 was targeted close to the transcriptional start site. The dOGT-dCas9 had virtually no effect on ^A^γ-globin mRNA expression at the −142, −182, −229, and −1034 sgRNA targeting sites. It did, however, have an activating effect on γ-globin mRNA expression at the −503 and −634 sgRNA target sites. When OGT-dCas9 was compared to dOGT-dCas9, OGT-dCas9 had statistically significant decreases in ^A^γ-globin mRNA expression at seven sgRNA target sites (−45, −142, −182, −229, −503, −634, and −1034). dOGT-dCas9 showed higher ^A^γ-globin mRNA repression at the −1 sgRNA target compared to OGT-dCas9 (**Figure 7a**). OGT-dCas9 showed a statistically significant decrease in ^A^γ-globin mRNA expression at both sgRNA target sites flanking the −566 GATA-1-FOG-1-NuRD repressor binding site (**Figure 7b**).

**Figure 7:**
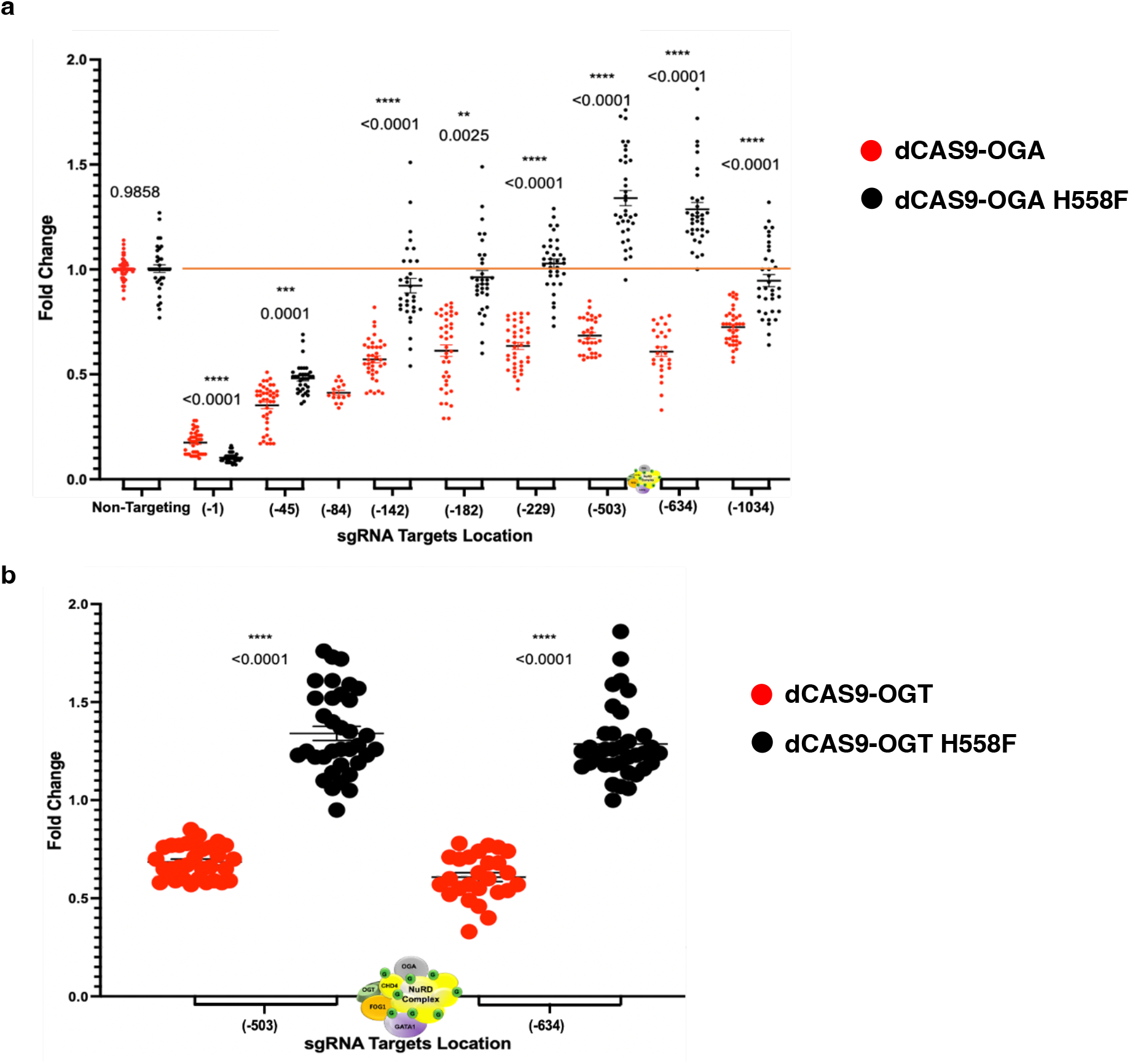
^A^γ-globin gene expression data in the presence of OGT-dCas9 or dOGT-dCas9 fusion constructs. (**a**) Relative mRNA expression of the ^A^γ-globin gene was determined by RT-qPCR. The OGT-dCas9 fusion protein represses transcription (red dots below the red line) of the ^A^γ-globin gene at all sgRNA target sequences when compared to non-targeting controls. The dOGT-dCas9 fusion protein had both repressive (−1 and −45) and activating (−503 and −634) effects on ^A^γ-globin gene expression (black dots above and below the red line) when compared to non-targeting controls. (**b**) OGT-dCas9 versus dOGT-dCas9 effect on ^A^γ-globin mRNA synthesis at sgRNA target sites flanking the −566 GATA-1-FOG-1-NuRD repressor binding site. Fold change was calculated using five housekeeping genes (B2M, RNA18sN5, PKG1, GAPDH, and YWHAZ). All samples were normalized to non-targeting controls. Data generated in at least five independent experiments are presented as mean + or − standard error; the two-tailed Student t-test statistic was applied with p < 0.05 considered to be a significant difference. The location of the −566 regulatory element is shown on the x-axis.

## Discussion

Previously, we demonstrated that O-GlcNAc plays a role in regulating human γ-globin gene transcription (30,31) suggesting that O-GlcNAcylation modulates the formation of a GATA-1-FOG-1-NuRD repressor complex that binds the −566/567 GATA-1 site of the ^A^γ and ^G^γ-globin promoters, respectively, when γ-globin genes are silenced (30,31). OGA inhibition using Thiamet-G (TMG) impairs the removal of O-GlcNAc from CHD4 and maintains repression of the γ-globin genes (30,31). In addition to CHD4, every protein of the NuRD complex is O-GlcNAcylated; however, the function of these modifications remains unknown (33,34). Thus, to investigate the ^A^γ-globin gene promoter for additional O-GlcNAc-regulated proteins bound at *cis*-regulatory elements and to further explore repression of the γ-globin genes via O-GlcNAcylation of the NuRD repressor complex at the −566 GATA-1 binding sites, we targeted our constructs using nine sgRNA sequences to the ^A^γ-globin promoter. We observed significant increases in ^A^γ-globin mRNA expression at the sgRNA-binding sites flanking the −566 GATA-1-FOG-1-NuRD repressor binding site when OGA-dCas9 was targeted and significant gene repression when OGT-dCas9 was targeted to identical sequences. Catalytically dead OGT activated γ-globin mRNA expression to a small extent at these sgRNA target sites. Our previous work showed that both OGT and OGA function at this location. In both cases, catalytically active enzymes produced the most profound effect on γ-globin mRNA expression at the −566 GATA-1 binding site.

In addition to the GATA-1-mediated recruitment of NuRD, this complex is also recruited to the γ-globin promoters at several additional *cis*-regulatory sequences by LRF/ZBTB7A and BCL11A (48). We observed statistically significant increases in γ-globin mRNA expression at the −229 sgRNA target site with both catalytically active and inactive OGA constructs. Proximal to this region, LRF/ZBTB7A and BCL11A recruit the NuRD repressor complex to silence the γ-globin gene expression. Catalytically active OGT had a significant repressive effect at the same sgRNA target site. Collectively, this data supports our original hypothesis that O-GlcNAcylation regulates γ-globin gene expression. Except for the sgRNA target site near the ^A^γ-globin transcription start site, O-GlcNAc manipulation had the most potent effects at known NuRD recruitment sites.

The OGT-dCas9 fusion protein had a significant repressive effect on ^A^γ-globin expression at all sgRNA target sequences analyzed; while the dOGT-dCas9 H558F fusion protein had both repressive and activating effects on ^A^γ-globin mRNA expression. Several possible mechanisms can explain the repressive effects observed with both OGT constructs near the transcriptional start site. First, OGT is known to interact with repressor complexes such as polycomb and the mSIN3A complex, potentially recruiting repressors to the site (7,8). Second, OGT modifies proteins within the pre-initiation complex (PIC) to promote assembly of the PIC with RNA Pol II. However, RNA Pol II activation might require OGA removal of O-GlcNAc from the C-terminal domain of RNA Pol II to initiate transcriptional elongation (12, 15). Third, the N-terminal domain of OGT might recruit repressive cofactors to the transcription start site since the catalytically dead OGT can repress, although not as well as the catalytically active construct (3). Fourth, the tetra-tri-co-peptide repeat domain (TPR) of OGT is hypothesized to act as a protein scaffold due to its similarity to importin-α and may be responsible for recruiting other proteins to *cis*-regulatory elements (59). The TPR of OGT may be recruiting endogenous OGT, thus explaining why dOGT had repressive effects on gene transcription at the −1 sgRNA target sequence (36). Moving forward, it will be essential to separate catalytic function from scaffold function for both enzymes. Lastly, a steric effect may disrupt endogenous regulatory elements when the constructs are targeted to the proximal ^A^γ-globin transcription start site (49). This does not seem likely, since the OGA-dCas9 is a larger fusion protein and had the opposite effect at those locations. However, this possibility cannot be completely ruled out, as OGA and OGT interact with different protein partners, which could have different steric effects at target sites.

When the OGA-dCas9 construct is targeted to the transcription start site, it activates transcription by removing O-GlcNAc moieties from the CTD of RNAP II, potentiating RNAP II release from the PIC by allowing RNAP II CTD phosphorylation (6,12–14). Mechanistically, O-GlcNAcylation and phosphorylation are mutually exclusive events, since they cannot simultaneously occur on the same amino acid residue. Thus, endogenous OGA occupancy at the γ-globin transcription start sites may increase the amount of RNAP II CTD phosphorylation leading to increased γ-globin expression (12). Mass spectroscopy interaction studies show that OGA does associate with Histone Acetyltransferases (HATs) (50,51). Possibly, some of the *cis*-regulatory elements that recruit OGA are not regulated by OGA catalytic function. For example, at the −1 sgRNA target site (transcription start site), OGA may affect gene expression by delivering a HAT to the transcriptional start site, possibly P300, a known interacting partner of OGA (51). Another possible explanation is that OGA functions as a homodimer at this site (52,53). Thus, we may be unintentionally directing endogenous catalytically active OGA to the γ-globin promoters via a dOGA-dCas9-OGA dimer. Increased endogenous OGA or increased phosphorylated RNAP II ChIP signals would confirm these mechanisms and support models that implicate O-GlcNAc in RNAP II function (6,12–14). OGT and OGA will form stable complexes, which might be involved in dynamically cycling the modification on and off proteins. (3,12,15,31,34,54–57). Thus, by targeting one enzyme, we may recruit the other enzyme increasing O-GlcNAc cycling rates without increasing or decreasing the overall O-GlcNAc levels at that target site.

To fully understand and make significant discoveries connecting gene regulation and O-GlcNAc signaling, we need to generalize beyond the γ-globin genes. Our CRISPR/dCas9 O-GlcNAc tools take advantage of the simple programmability of the CRISPR/Cas9 system to modulate O-GlcNAcylation at target sites. This new versatile and precise way to study the function of O-GlcNAcylation of proteins at *cis*-regulatory elements in the genome is a significant advance over existing approaches. The flexibility of our system allows it to easily be applied to genome-wide screens aimed at identifying O-GlcNAc-controlled *cis*-regulatory elements. Our system complements other recently described epigenetic editing tools and should help generate a better understanding of the function of O-GlcNAc in the genome. Furthermore, the tool will allow us to investigate the transcriptional roles O-GlcNAcylation plays in human diseases such as diabetes, cancer, Alzheimer’s disease, and Parkinson’s disease.

## Materials and Methods

### OGA and OGT-dCas9 Plasmid Synthesis

OGT and OGA dCas9 plasmids were synthesized using Gibson assembly (60). Briefly, the NEBuilder Assembly Tool (v1.12.18.) was employed to design the primers that were used to generate the PCR fragments comprising the final products. Plasmid pEF1α-FB-dCas9-puro (Addgene cat # 100547) contains a mammalian FLAG-tag, biotin-acceptor-site, and a dCas9 expression cassette. A pUC19 plasmid (Addgene cat # 50005) was utilized as template DNA to amplify the plasmid backbone fragment. Plasmids containing OGA (Origene cat # RC222411) or OGT (Origene cat # RC224481) cDNAs were used as template DNA for the PCR reactions in the Gibson assemblies. PCR reactions utilizing Q5 (New England Biolabs cat # M0492L), a high-fidelity DNA polymerase, were employed to amplify the needed DNA fragments. The correct PCR products were verified (fragment size by agarose gel electrophoresis) and purified from agarose gels using a Zymoclean gel DNA recovery kit (Zymogen cat # D4001). The resultant multiple overlapping DNA fragments were joined in a single isothermal reaction using a Gibson assembly cloning kit (New England Biolabs cat # E2611S). We designed our Gibson assembly primers to assemble the DNA fragments in the following order: 1) the elongation factor 1-alpha (EF1α) promoter fragment was attached to the pUC19 plasmid backbone; 2) the OGT or OGA fragments were attached to the EF1α promoter; 3) a NotI restriction site was added to the 3’ ends of the OGT or OGA sequences (C-terminus); 4) a dCas9 fragment was incorporated; 5) a FLAG-tag and biotin-acceptor-site was added to the 3’ end of dCas9 (C-terminus); and 6) the 3’ end of the FLAG-tag and biotin-acceptor-site fragments were annealed back to the pUC19 plasmid backbone, thereby circularizing the plasmid. The products of our Gibson assemblies were used to transform NEB 5-alpha competent E. coli cells (New England Biolabs cat # C2987H). Plasmids were isolated from individual bacterial colonies using a ZR plasmid miniprep kit (Zymo Research cat # D4054). The resultant plasmids were screened with restriction enzyme-digest and agarose gel electrophoresis. Sanger sequencing was used to verify that the potential positive clones were correctly assembled. Plasmid DNA maxi-preps (Millipore Sigma Maxiprep cat # NA0310-1KT) produced adequate amounts of high-quality DNA for downstream mammalian cell transfections.

### dOGA and dOGT-dCas9 Plasmid Synthesis

The OGT-dCas9 and OGA-dCas9 plasmid DNAs were then used to generate several of our experimental controls. These controls consisted of catalytically dead OGT and OGA dCas9 fusion constructs. To generate them, we perform site-directed mutagenesis. For OGT, we will introduce a H558>F mutation, and for OGA we introduce a D174>A mutation with a Q5 site-directed mutagenesis kit (New England Biolabs cat # E05545). These mutations produce stable mutant proteins that abolish the catalytic activity of the enzymes (38,39). Additionally, dCas9-p300 (Addgene cat # 83889) was ordered to serve as a positive control. As described above, we transformed competent cells and screen bacterial colonies for the correct plasmid constructs.

### Generation of OGA, dOGA, OGT, dOGT, and P300 -dCas9 stable cell lines

All constructs were linearized using PvuI restriction enzyme (New England Biolabs cat # R0150L). The linear products of these digestions were analyzed by agarose gel electrophoresis to confirm the full linearization of the plasmids. 50μg of linearized plasmid DNA was introduced into K562 cells via electroporation with Mirus Ingenio Electroporation Solution (cat # MIR50117) on a Bio-Rad GenePulser Xcell electroporation system according to manufactures protocol (2.5×10^6^ cells per 250ml of electroporation solution in a 4.0mm cuvette; General exponential decay pulse conditions with a voltage of 250V and capacitance of 1000mF were used). The K562 cells were maintained in Gibco Roswell Park Memorial Institute (RPMI) 1640 Medium (Corning Cat # 10-040-CV) supplemented with 10% heat-inactivated fetal bovine serum (Gemini cat # 100-106), 4mM L-glutamine (Corning cat # 25-005-Cl), 10mM HEPES (Corning cat # 25-060-Cl), 1mM sodium pyruvate (Corning cat # 25-000-Cl), 1x MEM Nonessential Amino Acids (Corning cat # 25-025-Cl), and 100U/ml penicillin and 100μg/ml streptomycin (Corning cat # 30-002-Cl). The cells were incubated at 37°C in a humidified 5% CO_2_-containing atmosphere for 72 hours. The cells were then treated with 2μg/ml of puromycin (InvivoGen cat # ant-pr-1). After 72 hours, limiting dilution was utilized to generate single clone lines. Clonal cell lines that contain similar gene cassette copy numbers and fusion protein expression levels were selected for subsequent experiments. The following specific fusion protein cell lines were generated: positive control dCas9-p300, OGT-dCas9, OGA-dCas9, and catalytically dead versions of dOGT-dCas9 and dOGA-dCas9.

### Genotyping of Stable Cell Lines

Genotyping was done by DNA PCR using Q5 (New England Biolabs cat # M0492L) and confirmed the presence of the expression cassettes.

### Western Blots

Cells were lysed on ice as described previously (34). 50μg of whole-cell lysates were used for electrophoresis and subjected to Western blots as previously described (34). Western blots revealed the presence of the fusion protein constructs with antibodies specific to Cas9 (Abcam cat # ab204448), OGT, OGA, and FLAG (Abcam cat # ab1162). GAPDH was utilized as a load control (Abcam cat # ab9484). Anti-OGT (AL-34) and OGA (345) were gracious gifts from Dr. Gerald Hart at the Johns Hopkins University School of Medicine. Primary antibodies and secondary antibodies for immunoblotting were used at 0.5 μg/ml and 1:10,000 dilution, respectively. The Anti-Cas9 antibody was used in accordance with manufacturer’s protocol.

### Synthesis of sgRNAs Plasmids

In order to target our fusion protein constructs, we designed and synthesized sgRNA expression cassettes as previously described (61,62). Briefly, CRISPR targets were selected using the ENCODE database. sgRNA oligos were designed to have a BsmBI complementary overhang so that they could be cloned into the LentiCRISPRv2 hygro plasmid (Addgene cat # 98291). The sgRNA oligos were ordered from Integrated DNA Technologies IDT. Complimentary sgRNA oligos were phosphorylated with T4 polynucleotide kinase (New England Biolabs cat # M0201S) according to manufactures protocol and annealed to each other using a Bio-Rad C1000 thermocycler with the following reaction conditions: 1) 37°C for 30 minutes; 2) 95°C with a ramp down to 25°C at 5°C/minute. Next, 50μg of the LentiCRISPRv2 hygro plasmid was digested with BsmBI (ThermoFisher cat # FD0454) and dephosphorylated with FastAP Thermosensitive Alkaline Phosphatase (ThermoFisher cat # EF0654) according to manufactures protocol. After restriction enzyme digestion and dephosphorylation, the LentiCRISPRv2 hygro plasmid was purified using a 0.7% agarose gel and a Zymoclean gel DNA recovery kit (Zymo Research cat # D4001). The prepared sgRNA oligo duplexes were then ligated into the purified LentiCRISPRv2 hygro plasmid. The products of these reactions were used to transform NEB Stable competent E. coli cells (New England Biolabs cat # C3040H). Plasmids were isolated from individual bacterial colonies using a ZR plasmid miniprep kit (Zymo Research cat # D4054). The resultant plasmids were screened with restriction enzyme-digest and agarose gel electrophoresis. Sanger sequencing confirmed that the sgRNAs were correctly ligated into the LentiCRISPRv2 hygro plasmid backbone. Plasmid DNA maxi-preps (Millipore Sigma Maxiprep cat # NA0310-1KT) produced adequate amounts of high-quality DNA for downstream mammalian cell transfections. The sgRNA sequences used in our experiments are listed in (**Table 1)**.

**Table 1.**
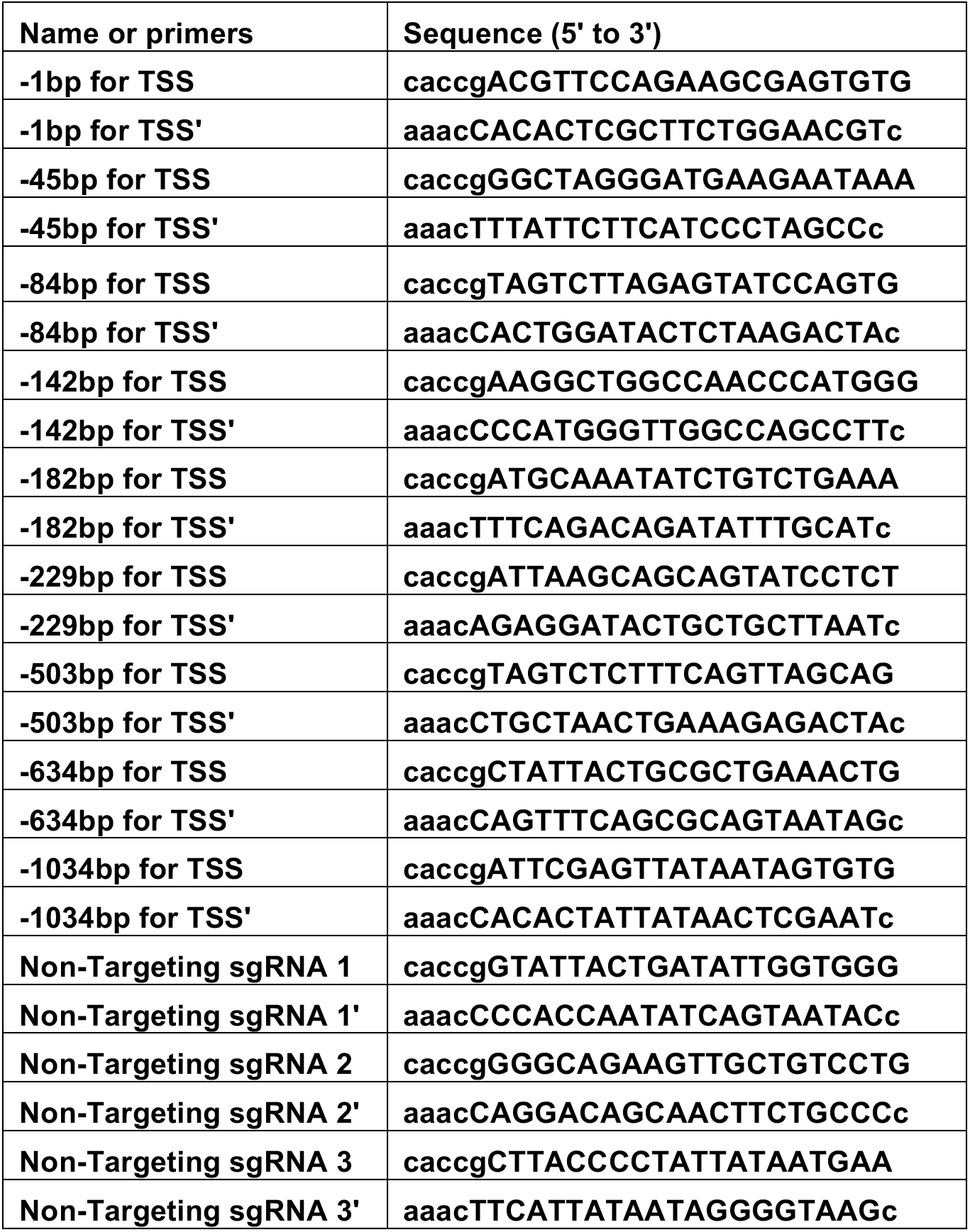
sgRNA Sequences.

### Generation of sgRNA Stable Cell Lines

All constructs were linearized using FspI restriction enzyme (New England Biolabs cat # R0135S). The linear products of these digestions were analyzed by agarose gel electrophoresis to confirm plasmid linearization. 50μg of linearized plasmid DNA was introduced into K562 parent monoclonal cell lines (OGA, dOGA, OGT, dOGT, and P300-dCas9) as described above. After 72 hours, 400μg/ml of Hygromycin (InvivoGen cat # ant-hg-1) was added to the cultures to select for positive sgRNA expressing cells. RT-qPCR was used to determine sgRNA expression levels. Stable cell lines expressing similar levels of sgRNAs were used in subsequent experiments.

### Total RNA Isolation for RT-PCR

Total RNA was isolated from 5 × 10^6^ cells using TRI Reagent (Sigma cat # T9424) according to the manufacturer’s instructions. For the reverse transcriptase (RT) reactions, 0.5 μg of total RNA was reverse-transcribed to cDNA using iScript Reverse Transcription Supermix (Bio-Rad cat # 170-8841) following the manufacturer’s instructions. Reactions were incubated in a thermal cycler (Model 2720, Applied Biosystems) using the following protocol: priming 5 min at 25°C, RT 20 min at 46°C, RT inactivation 1 min at 95°C, and hold at 4°C.

### Reverse Transcription Quantitative Polymerase Chain Reaction (RT-qPCR)

RT-qPCR was performed as described previously described and adhere to the MIQE guild lines (34). The primer sequences for measuring target gene transcription levels are listed in (**Table 2)**. The reactions were run in a CFX96 Touch Real-Time PCR Detection System (185-5195, Bio-Rad). All experiments have been performed a minimum of five times (n > 5)

**Table 2.**
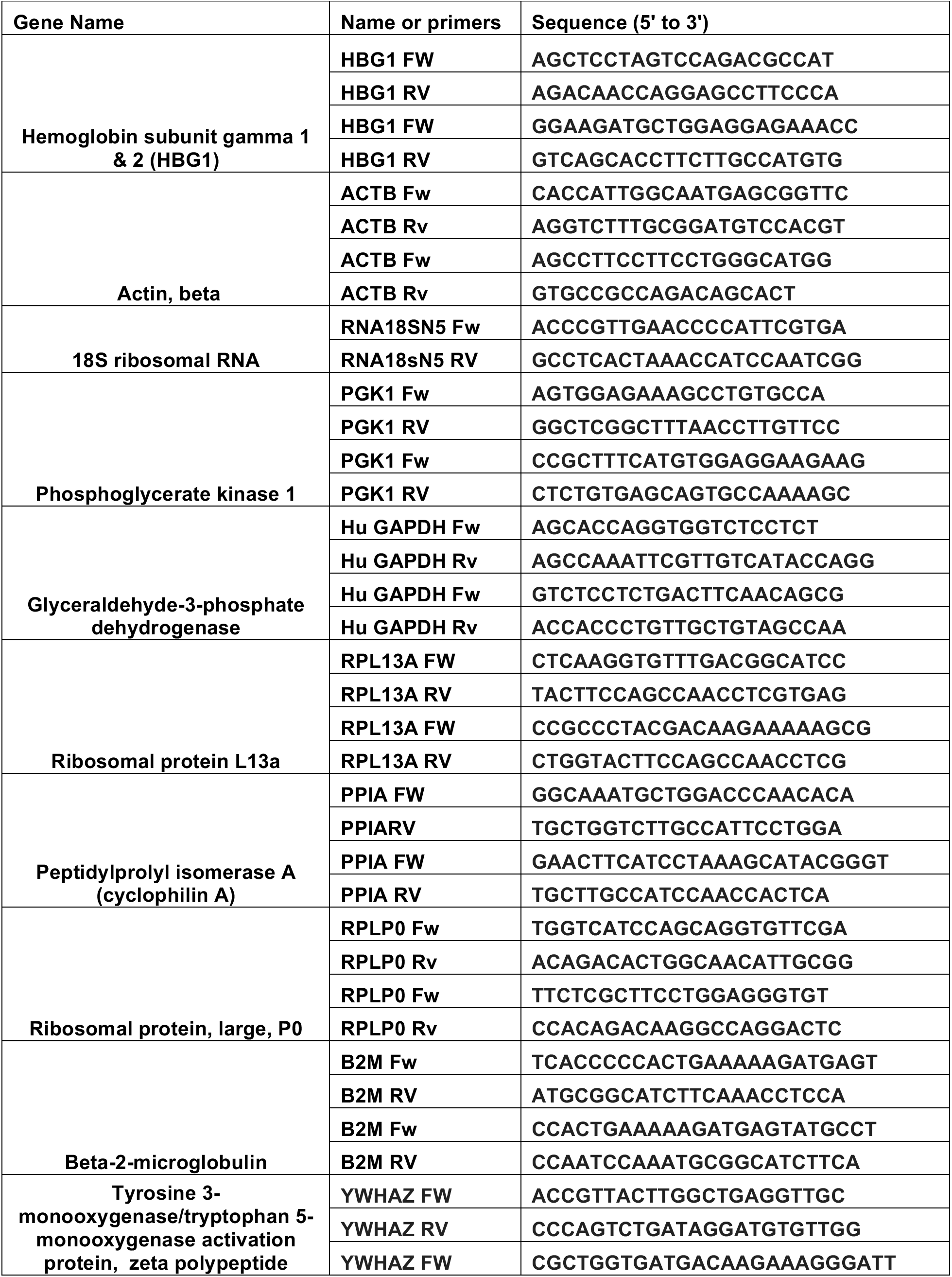
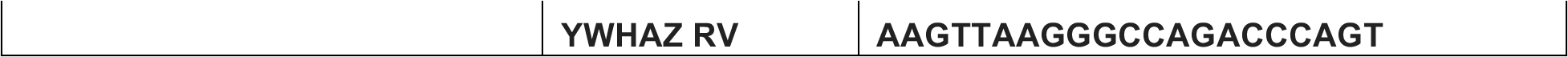
qPCR primers.

### qPCR Data Analysis

Quantification cycle (Cq) values were calculated by CFX Manager™ software (Bio-Rad). For cDNA RT-qPCR data, the dynamic ranges of RT and amplification efficiency were evaluated before applying the ΔΔCq method to calculate relative gene expression change. The transcription level of the target gene was normalized to the internal control as fold change. Data generated in at least five independent experiments are presented as mean + or − standard error; the two-tailed Student t-test statistic was applied with p <0.05 considered to be a significant difference.

## Notes

### Competing Interest Statement

The authors have declared no competing interest.

